# Effects of productive management of an agroforestry system on the associated diversity of parasitoids (Hymenoptera: Braconidae)

**DOI:** 10.1101/2021.10.01.462815

**Authors:** Cecilia Marisol Pech Cutis, Luis Enrique Castillo Sánchez, Jorge Rodolfo Canul Solís, Ermilo López Coba, Nery Maria Ruz Febles, Maria Jose Campos-Navarrete

## Abstract

Tropical agroecosystems have emerged from the continuous modification of natural environments, as a sustainability alternative for food production and biodiversity conservation. This work explores how the diversity of parasitoids is modified in environments where plant diversity is limited e.g. crops and when these are adjacent to secondary vegetation, i.e. a scenario fragmented continuously in a limited space. It was found that there is no direct effect of plant diversity in the group of parasitoids studied; but the number of specialist species is high, which indicates that in diversified agroecosystems these probably function as remnants of natural habitat or as a refuge for parasitoids that disperse to different types of management within the agroecosystem. Therefore, it is necessary to consider in future studies the controls exerted by the plant diversity effects bottom-up and consumer top-down. Adding to this the context of the interactions that occur in agroecosystems.

## Introduction

Agroecosystems arise in the tropics derived from the loss of vegetation cover, the conversion and fragmentation of natural areas in ecosystems. These activities modify the species that inhabit and depend on vegetation,with effects that depend on the range of the species and its habitat requirements (Scolozzi & Geneletti, 2012; Campos-Navarrete et al 2015a). On the other hand, the speed in the loss of species has increased the interest in the study of biological diversity in agroecosystems, mainly insects, because they constitute the most important fraction of the diversity of a territory since they provide multiple ecological services to ecosystems (Kleinet al 2002).

Parasitoids are known for their high number of species and their habits that result in the provision of regulatory services for the populations of their hosts (Lepidoptera, Coleoptera and Diptera) (Shaw & Huddleston, 1991; Abdala-Roberts et al 2016a). In agroecosystems, the study of these organisms is interesting from the anthropogenic point of view (integrated pest control) and even little known in terms of the mechanisms that regulate it (Nicholls, 2008; Schmidt et al 2003; Thies et al 2005). Also in agroecosystems, plant species diversity has effects on secondary productivity comparable to natural systems. In this sense, it has been observed that environments with greater diversity of plant species promote increases in richness and abundance in trophic levels (Abdala-Roberts et al 2016 b; Castillo-Sánchez et al 2019).

The effect of land use, in conjunction with the associated plant diversity at higher trophic levels, has been explained primarily because greater diversity, by generating a more complex environment, consequently offers a greater number of shelters and prey (Russell, 1989; Obermaier, et al 2008; Moreira et al 2016),which in turn generates increases in predation rates, causing a reduction in the abundance of parasitoid prey (“up-down” effects). Evidently, this type of effects are highly relevant to be considered in the design of productive systems such as forestcrops, due to their potential in pest control (Russell, 1989; Abdala-Roberts et al 2015; 2016a). This last factor can reduce or increase the abundance of prey (herbivores) depending on characteristics such as the specialization of their diet (generalists vs. specialists) and their interactions (Campos-Navarrete et al 2015b). For example, there is evidence that for specialists the effects of an increase in diversity can be negative, due to the low density of their priority resource (Hambäck et al 2014).

In contrast, for generalists the effects of increased diversity can vary and, in some cases, can be positive due to their mixed diet and increased availability of places of refuge (Unsicker et al 2008; Castagneyrol et al 2013). In this sense, the presence of herbivores can mediate their interaction with the next trophic level of consumers where parasitoids are included (Abdala-Roberts et al 2016b).

The present work explores the effect of four productive areas in an agroecosystem of multiple production in relation to the diversity of parasitoids (Braconidae) associated. Particularly changes in richness, abundance, similarity in areas and in parasitism strategies represented by the proportion of koinobiont (specialist) and idiobiont (generalist) species. Trying to infer how the diversity of parasitoids is modified in environments where plant diversity is limited e.g. crops and when these are adjacent to secondary vegetation, i.e. a scenario with continuously fragmented in a limited space.

## Materials and Methods

### Study area

The present work was carried out in the east of Yucatán in the municipality of Tizimin, Yucatán, Mexico. This area is characterized by the conversion of land use from native jungle to grasslands for the production of pasture for cattle feed. Livestock represents 30% of the economic activity of this area (INEGI 2015). The agroecosystem of study is located in the area of agricultural and livestock production of the TecNM Campus Tizimin, located at the end of the airport Cupul s/n C.P. 97700 with the coordinates 21°09′29” N 88°10′21”W.

The agroecosystem has three areas of crops: a plantation of *Cocos nucifera* “PC” (monoculture),a plantation of Citrus *lemon* “PL” (monoculture), in the livestock production area is located the grassland area with star grass *Cynodon sp*. named grassland “PT” and the fourth type is the matrix of secondary vegetation with more than 30 years of recovery “VS”. This contrast in a limited space provides a frequent scenario today, originated by the processes of fragmentation, originating contrasting sites.

### Fieldwork

For the capture of insects were used malaise traps, which capture large numbers of organisms, widely recommended and used to capture parasitic Hymenoptera (Noyes, 1982) (Nieves-Aldrey & Castillo., 1991). The orientation of the trap was from north to south, because it is more effective if the openings are placed in the position where the wind comes from. For the placement of the trap it was considered that it was far from the edges of each area, in order to reduce the edge effect and between traps there was a distance of 500 m, to ensure the independence of the samples. The traps remained active every day during the period from October 2015 to March 2016, were checked every fortnight and the specimens placed in jars with 70% alcohol.

### Identification of the specimens

The samples were identified in the Laboratory of Parasitology of the TecNM Campus Tizimín, before the identification the braconids were separated from the rest of the insects, later they were counted and labeled with the corresponding data, for their identification the specimens were assembled according to the technique traditionally used for these organisms. Identification was carried out using the taxonomic key for Braconidae (Wharton et al 1998).

The assembly was carried out using entomological pins correctly placing each insect, which were also labeled for a systematic control of these. The reason for mounting is that in this way it is easier to observe insects when they are dry, since when they are wet it is difficult to observe certain characteristics.

The material was determined up to the taxonomic level of genus subsequently the concept of morphospecie was used in which (Simpson, 1961; Mayr & Ashlock., 1991; Delfín & Burgos 2000) mention that due to the existing difficulty in determining parasitoids at a specific level it is advisable to use this concept to identify and separate individuals who present different morphological characteristics. In addition, morphospecies were classified according to their parasitism strategies as koinobiont species (specialists) and idiobionts (generalists), with the help of specialized literature (Goulet & Huber, 1993). All the collected material is sheltered in the TecNM Campus Tizimín.

### Data analysis

We performed a classical diversity analysis in the braconid community in all four areas using the Shannon-Wiener diversity index. This index expresses the uniformity of the values of importance across all species in the sample; assumes that all individuals are randomly selected, and that all species are represented in the sample (Ludwing and Reynolds, 1988; Magurran, 1988).

The similarity between the braconid communities was estimated using the Sorensen index (Krebs, 1989) which is based on the presence/absence of species and the number of species whether common or rare (Spellerberg, 1991). It relates the number of species in common to the arithmetic mean of the species at the sites (Magurran, 1988).

To estimate the representativeness of the species richness of the samples in each type of vegetation, the EstimateS version 9 program was used, which presents specialized estimators for different types and sizes of data. We used ice wealth estimators based on the number of rare species (those observed in less than 10 sampling units) and jacknife first order which is based on the number of unique species (Colwell and Coddington, 1994).

With the data of the morphospecies recorded by area, using relative abundance, as the response variable, the change between the four areas was evaluated. This is through a model that explores the quantitative response of braconids to areas. This model used as the main factor the area with four levels (PC, PL, PT, VS). The second model explored the parasitism strategies of braconist species, using as the main factor parasitism strategy with two levels (Idiobionts “I” and Koinobiontes, “K”), in relation to abundance. The third model explored the interaction between the two main factors mentioned above in the relative abundance of braconids. We fitted all models using the Penalized Quasi-Likelihood method (Crawley 2007). We conducted GLMM analyses using the R statistical package v. 3.01. (R Core Team 2020). We used a posteriori contrasts to test for differences among pairs of means for a given factor within each level of the other factor (Crawley 2007).

## results

### Collected specimens

A total of 2031 specimens of braconids belonging to 20 subfamilies, 47 genera and 140 morphospecies (Apendice 1) were collected. In general, the subfamily Microgastrinae was the most abundant representing 49% of the total number of individuals captured, other abundant subfamilies were Doryctinae and Hormiinae with 8% and 6% individuals respectively, the least abundant subfamilies were Ichneutinae and Macrocentrinae which are represented by an individual equivalent to 0.05% (Table 1).

**Table 1.** Number of genera (G), morphospecies (S) and abundance (A) by subfamilies in each area PC= Coconut Plantation, PL= Lemon Plantation, PT= Pastizal, VS=Secondary Vegetation and the Total, in the agroecosystem.

| subfamily | PC |  |  | pl |  |  | Pt |  |  | vs |  |  |  |  |
| --- | --- | --- | --- | --- | --- | --- | --- | --- | --- | --- | --- | --- | --- | --- |
| Number of individuals | 447 |  |  | 1232 |  |  | 194 |  |  | 158 |  |  | total |  |
|  | G | s | to | G | s | to | G | s | to | G | s | to | to | s |
| **Agathidinae** | 2 | 5 | 32 | 2 | 4 | 24 | 3 | 5 | 6 | 2 | 2 | 4 | 66 | 8 |
| **Alysiinae** | 1 | 1 | 3 | 4 | 4 | 33 | 2 | 2 | 2 | 2 | 2 | 5 | 43 | 5 |
| **Aphidiinae** | - | - | - | - | 1 | 78 | - | - | - | - | - | - | 78 | 1 |
| **Blacinae** | - | - | - | - | - | - | - | - | - | 1 | 2 | 7 | 7 | 2 |
| **Braconinae** | 3 | 5 | 44 | 4 | 6 | 39 | 3 | 5 | 7 | 1 | 1 | 3 | 93 | 7 |
| **Cardiochilinae** | - | - | - | 1 | 2 | 18 | - | 1 | 1 | - | 1 | 1 | 20 | 2 |
| **Cheloninae** | 2 | 6 | 11 | 3 | 9 | 63 | 2 | 5 | 9 | 2 | 3 | 5 | 88 | 16 |
| **Doryctinae** | 4 | 1<br>0 | 19 | 6 | 24 | 95 | 3 | 9 | 25 | 2 | 1<br>4 | 24 | 163 | 31 |
| **Euphorinae** | 3 | 3 | 4 | 3 | 4 | 25 | 1 | 1 | 2 | 1 | 1 | 2 | 33 | 6 |
| **Homolobinae** | - | - | - | 2 | 2 | 6 | 2 | 2 | 2 | - | - | - | 8 | 3 |
| **Hormiinae** | 4 | 7 | 17 | 7 | 15 | 84 | 2 | 2 | 8 | 5 | 7 | 11 | 120 | 16 |
| **Ichneutinae** | - | - | - | 1 | 1 | 1 | - | - | - | - | - | - | 1 | 1 |
| **Macrocentrinae** | 1 | 1 | 1 | - | - | - | - | - | - | - | - | - | 1 | 1 |
| **Mendesellinae** | 1 | 1 | 2 | 1 | 1 | 7 | - | - | - | - | - | - | 9 | 1 |
| **Meteorinae** | 1 | 1 | 1 | - | - | - | 1 | 1 | 1 | - | - | - | 2 | 1 |
| **Microgastrinae** | - | 1<br>8 | 24<br>7 | - | 19 | 572 | - | 1<br>9 | 12<br>1 | - | 1<br>0 | 46 | 986 | 23 |
| **Miracinae** | 1 | 1 | 4 | 1 | 1 | 17 | - | - |  | 1 | 1 | 6 | 27 | 1 |
| **Opiinae** | 1 | 5 | 47 | 1 | 7 | 143 | 1 | 4 | 6 | 1 | 4 | 39 | 35 | 7 |
| **Orgilinae** | 2 | 2 | 5 | 1 | 1 | 4 | 1 | 2 | 4 | 2 | 2 | 4 | 17 | 4 |
| **Rogadinae** | 2 | 2 | 10 | 2 | 3 | 23 | - | - | - | 1 | 1 | 1 | 34 | 4 |
| **total** | **28** | **68** | **447** | **39** | **104** | **1232** | **21** | **58** | **194** | **21** | **51** | **158** | **2031** | **140** |

### Richness and abundance of species by typearea

Wealth in each area of the agroecosystem was distributed as follows from highest to lowest. PL with the largest number of subfamilies, followed by PC, VS and PT (Table 1). As for gender from highest to lowest PL, PC, VS and PT (Table 1). The morphospecies were distributed from highest to lowest in the areass in the following order first PL, second PC and with the same number PT and VS (Table 1). Finally, for the number of individuals the largest was concentrated in PL, followed by PC, PT and finally VS (Table 1). For wealth in general it was observed that the PL and PC,considereds monocultures concentrated the largest number of individuals and taxonomic wealth, while VS occupied the third site and with the least wealth and abundance the PT with the simplest vegetation structure composed solely of grass.

The general pattern for the number of rare and unique morphospecies, as well as species diversity according to the Shannon Index by area is described below. For rare species the order of greatest number was in PT, PL, PC and VS (Table 2). As for unique species from highest to lowest these were found in PC, PT, PL and finally VS (Table 2). In general, it is observed that the VS considered a polyculture and with greater complexity was the one with the lowest number of rare and unique species present. This contrasts with what was found for the Shannon-Wienner diversity index, where the highest diversity was observed in PL, followed by VS, finally PC and PT (Table 2).

**Table 2.** Shannon-Wiener diversity index, Total number of morphospecies, Singletons, Doubletons, Unique species and rare species in PC coconut plantation areas; PL lemon plantation; PT grassland and VS secondary vegetation in the agroecosystem.

|  | *PC* | *pl* | *Pt* | *vs* |
| --- | --- | --- | --- | --- |
| ***Shannon-Wiener index*** | 3.43 | 3.77 | 3.41 | 3.45 |
| ***Total number of species*** | 68 | 104 | 58 | 51 |
| ***Singletons*** | 28 | 25 | 28 | 20 |
| ***Doubletons*** | 7 | 13 | 12 | 14 |
| ***Unique species*** | 35 | 31 | 32 | 30 |
| ***Rare species*** | 35 | 38 | 40 | 34 |

### Specific wealth estimation

The wealth estimators used were ICE and Jacknife of the first order. According to the ICE, 60% of the species were captured on average for the agroecosystem, while for the jacknife of the first order 70% were captured according to the estimate for the agroecosystem. Table 3 shows in particular the percentage of wealth estimators by area.

**Table 3.**
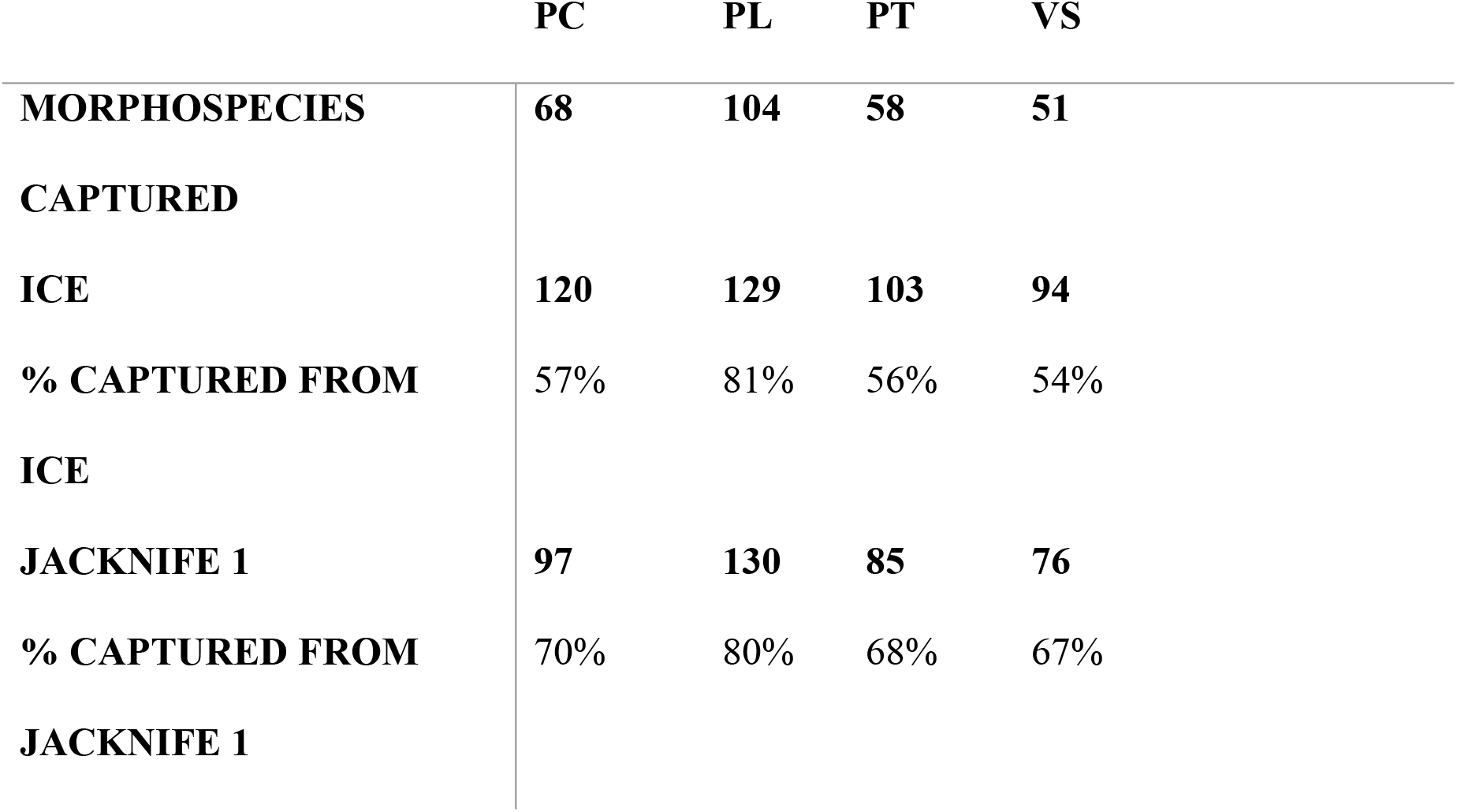
Ice and Jacknife wealth estimators of the first order (Jacknife 1),with respect to the morphospecies captured, calculated in the areas of PC coconut plantation; PL lemon plantation; PT grassland and VS secondary vegetation in the agroecosystem.

### Similarity of communities

The Sorensen index indicated that the sites that presented the greatest similarity were PC with VS, followed by PT with VS, in third place, PL with VS and those that presented the least similarity were PC with PT (Table 4). The above highlights the importance of conserving remnants of secondary vegetation in agroecosystems, since it was observed that VS is similar to crop areas.

**Table 4.** Similarity of the braconid community in the areas of PC coconut plantation; PL lemon plantation; PT grassland and VS secondary vegetation in the agroecosystem.

|  | PC | PL | PT | VS |
| --- | --- | --- | --- | --- |
| PC | - | 0.41 | 0.33 | 0.65 |
| PL | - | - | 0.49 | 0.54 |
| PT | - | - | - | 0.58 |
| VS | - | - | - | - |

### Effect of the areas on the abundance

As for the quantitative response in the abundance relative to the area of cultivation and the functional role of the braconist in this system, it was observed that the Area has a statistically significant effect on abundance (F _1, 3_ = 2.8125 p *=* 0.000). Higher values of relative abundance were observed in PC, PT and VS with respect to PL (Figure 1). As for the parasitism strategy, a significant effect was found on abundance (F_1.2_=8.5947 p *=*0.000), being greater in koinobionts (specialists) compared to idiobionts (generalists) (Figure 2). However, no interaction effect was found between the Area and parasitism strategies (F_1.1_= 0.6097 *p* = 0.*65*)inthis agroecosystem.

**Figure 1.**
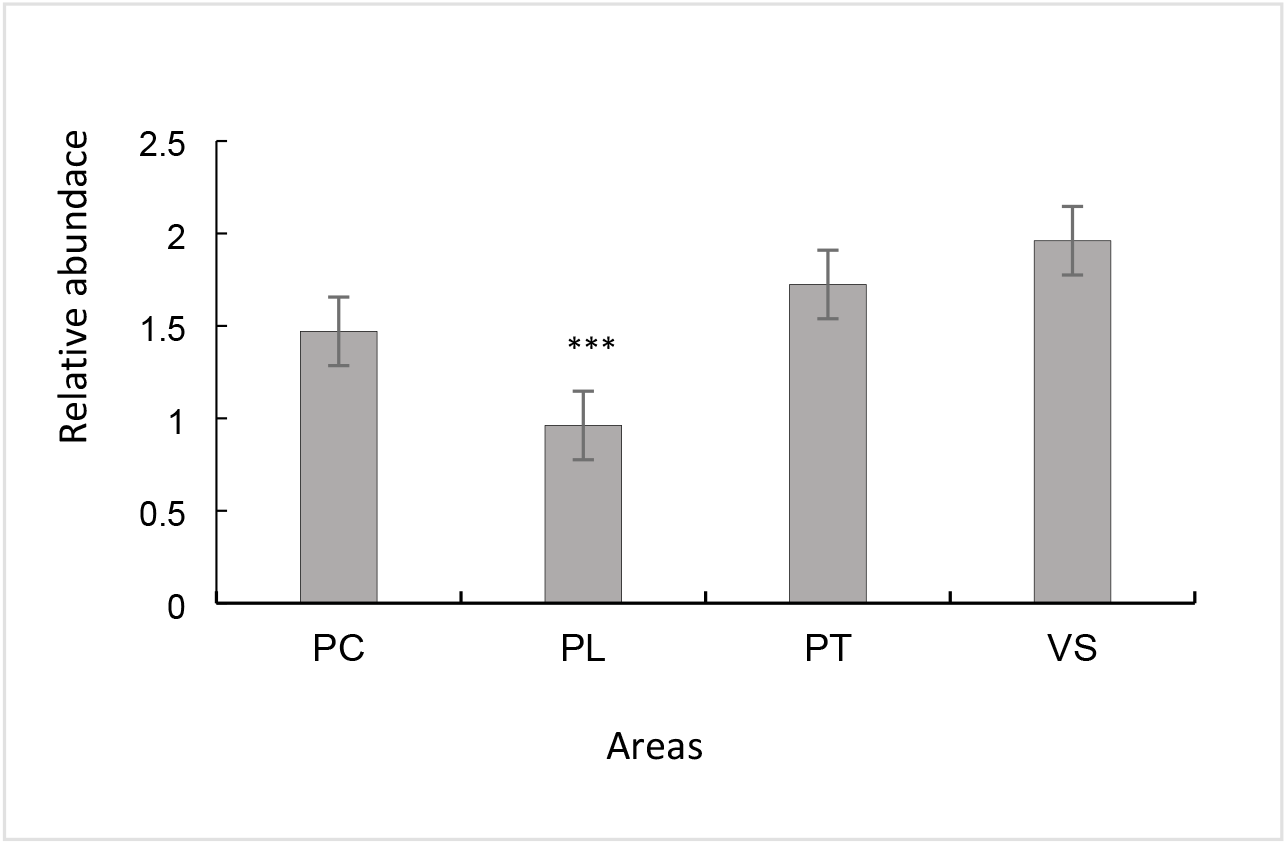
Average relative abundance (EE±) of braconid morphospecies per pc coconut plantation area; PL lemon plantation; PT grassland and VS secondary vegetation in the agroecosystem (***p<0.05)

**Figure 2.**
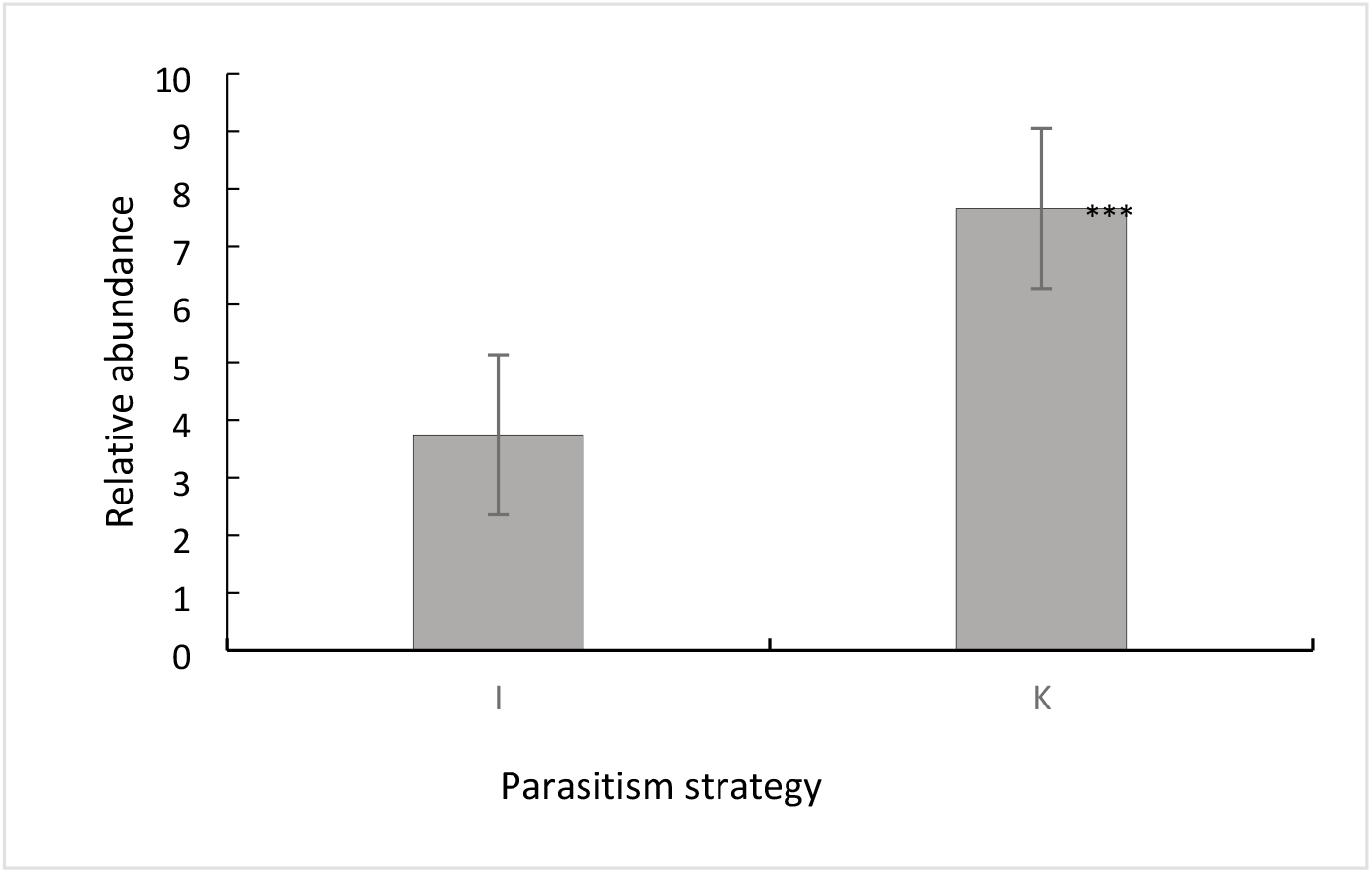
Relative average abundance by parasitism strategy (EE ±) of braconid morphospecies per area PC coconut plantation; PL lemon plantation; PT grassland and VS secondary vegetation in the agroecosystem (***p<0.05)

### Proportion of the estrategia of parasitism

The distribution was as **follows**, in general the Koinobionte (specialist) represented 63% on average, while 36% on average is classified as Idiobiont (generalists) and only 1% that corresponds to the registror a species of the genus Epsilogaster (Mendesellinae) of which its biology is unknown (Table 5). The percentage from highest to lowest in the areas of koinobiont species was higher percentage was presentedor in PT, followed by PC, VS and finally PL (Table 5). As for Idiobionts with the highest and equal percentages PL and VS, followed by PC and finally PT (Table 5). This result suggests that in a diversified agroecosystem, species of braconide specialist nests are more abundant, mainly in complex areas such as monocultures; and generalists concentrate on areas of greater complexity such as secondary vegetation.

**Table 5.**
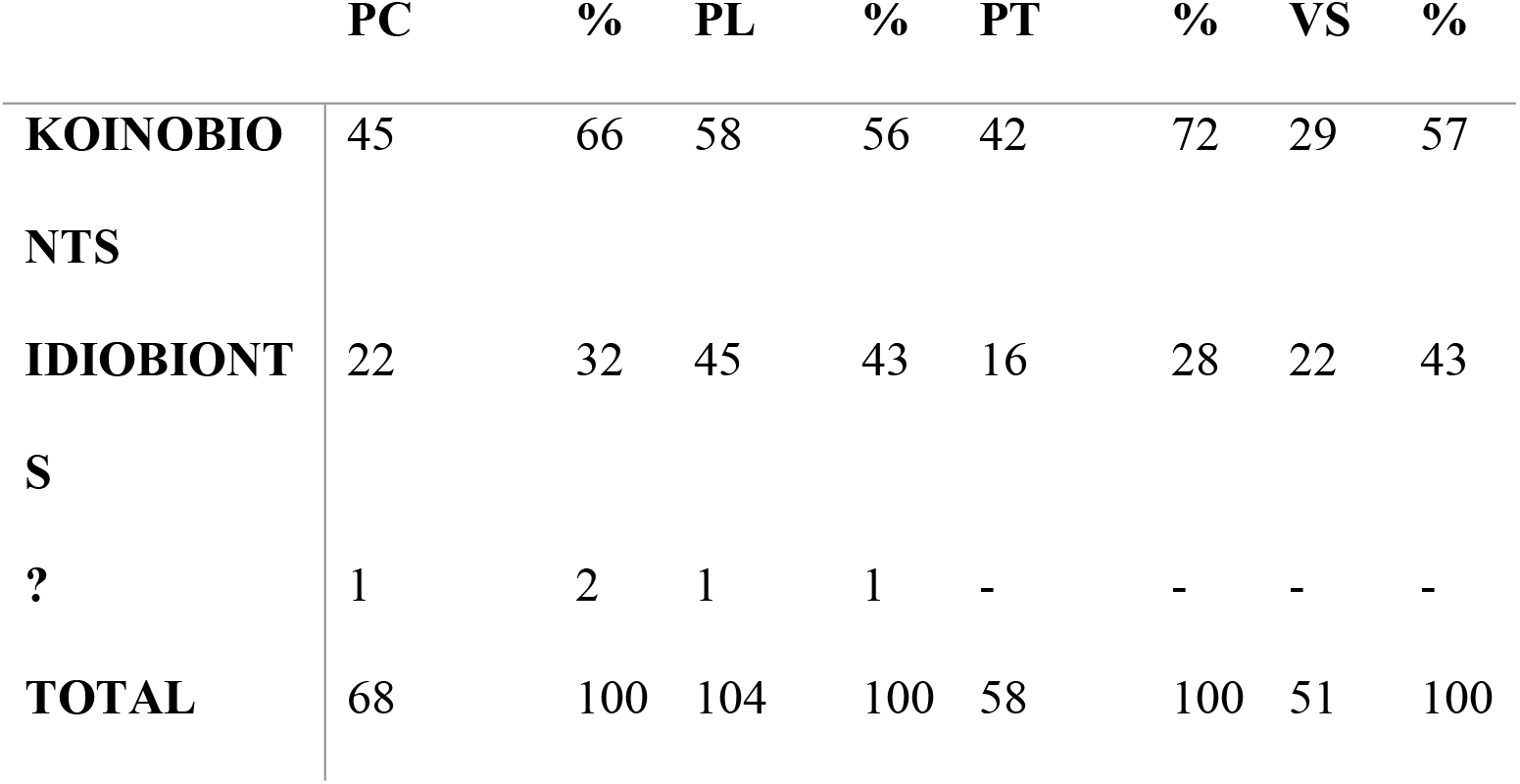
Strategies of parasitism and morphospecies in which it is not known (?=Unknow) expressed in total and percentage for PC coconut plantation; PL lemon plantation; PT grassland and VS secondary vegetation in the agroecosystem.

## Discussion

Braconids are generally considered to be of great importance for their participation in the natural control of other insects and for their use in biological control programs for forest pests, fruit trees, vegetables and extensive crops worldwide (Coronado et al 2010). But the mechanisms of this possible regulation in management systems are little explored, particularly in agroecosystems that expect high levels of plant diversity and maintain high levels ofbiodiversity, with abiotic and abiotic changes that can affect insect communities (Klein et al 2002).

In the study of this diversified agroecosystem, it was found that the areas contain high levels of diversity of bracornests, with the presence of high percentages of specialistsin the crops and communities of these organisms very similar to the remaining secondary vegetation. This supports the benefit of agroecosystems as conservation strategies, as some insects (e.g. eumenid wasp and solitary bees) do not decline significantly to the modification of the environment (Klein et al 2002),which apparently happens in the case of bracornests.

According to the data obtained in this work it was observed that Microgastrinae was the most abundant subfamily, which is expected. For this subfamily it has been estimated between 4000 to 10 000 species inthe world (Joneset al 2009). The high abundance of microgastrinae may be due to the fact that they are braconids with a wide range of hosts, which attack almost all families of Lepidoptera, except the family Hepialidae (Shaw, 1994). And it includes species of small size and with dark colors that possibly help them not to be seen by their predators (Rathcke and Price, 1976; Hawkins et al 1992). The subfamilies Ichneutinae and Macrocentrinae were the least abundant, this is in line with the predicted for these subfamilies which estimate there are between 100 -300 species worldwide (Jones et al 2009).

The study of braconids in Mexico is focused on three areas and they are, the knowledge of their taxonomic richness, which includes both faunal studies and descriptions of new taxa; research in ecology, mainly using these organisms as indicators of biodiversity; their use as biological control agents of other insects, with potential applications in agriculture and forestry activities (Coronado et al 2010; Coronado, 2011).

Recent studies have reported 318 genera of Braconidae for Mexico, 194 belonging to Yucatán (Coronado and Zaldívar, 2014). In this work, 47 genera were registered, which constitutes 24% of what was recorded for Yucatán. This family is well represented in the peninsula as it is the state with the highest number of genera recorded so far. It should be mentioned that the high number of genera found for this state is due to the fact that it is very close to the Gulf of Mexico and because it is one of the states where national and foreign specialists work and /or where more collections have been made (Coronado and Zaldívar, 2014).

With the data obtained in this agroecosystem it was possible to observethat the areas of monocultures (e.g. PL), presented greater diversity of species, in terms of taxonomic richness and abundance of individuals. However, for parasitoid hymenoptera considered “specialised enemies” are not affected by the manipulation of plant diversity (Koricheva et al 2000; Abdala-Roberts et al 2016a). This was observed in three of the four areas where there were no significant differences in the abundance of bracor nests.

In this agroecosystem it was found that the largest number of individuals were classified as specialists (koinobionts) and that the presence of these is independent of the area of cultivation. One mechanism that could explain the patterns found in this agroecosystem may be More specialization (sensu Obermair et al 2008)which indicates specialization prevents competition, so productive communities support a greater number of species.

In this agroecosystem it remains to explore the role of the abundance of herbivores, which has been pointed out in other cases, as a factor that suggests that the presence ofherbivores affects the number of parasitoids and the density of their population with respect to landscape, climate and management (Koricheva et al 2000; Schmidt et al 2003). This indicates that the presence of herbivores in crops, causedby the concentration of a single resource would affect the incidence of certain pests, which result in an important source for the attraction of certain parasitoids, increased consumer abundance as proposed by the More Individuals Hypothesis (Srivastava & Lawton 1998).

On the other hand, polyculture (VS) can present greater plant diversity than other sites, probably functioning as a remnant of natural habitat or as a refuge for parasitoids that disperse to different types of management within the agroecosystem, depending closely on the management (Thies and Tscharntke, 1999) which should be explored.

The results correspond to what was reported by Askew and Shaw (1986), Hawkins (1994), Ruiz-Guerra et al (2014), Rodríguez-Soliset al(2016) where they indicate that the braconid community is dominated by koinobiont parasitoids, mentions that the specialty of the host is not related to the degree of disturbance and therefore koinobiont parasitoids can occur both in the early stage and in the late succession phase of vegetation.

Finally, the controls exerted by the plant diversity bottom-up and consumer top-down effects should be considered in future experimental work. Adding to this the context of the interactions that occur in agroecosystems must be studied with respect to the communities (e.g. insects, animals) associated with the different trophic levels and the functional role they take in these systems.

## Acknowledgments

TecNM (10396.21-P; 6581-18P; 6194.17P).

## Appendices

### Appendix 1

List of subfamilies, morphospeies, abundances and biology of braconids in four sites: PS=Parasitims strategy, *K* =Koinobionte, I = Idiobionte, ?= Unknow, PC = Coconut Plantation, PL = Lemon Plantation, PT = P astizal, *VS = Secondary Vegetation*and TL = Total *in the* agroecosystem.

| subfamily | species | B | PC | pl | Pt | vs | Tl |
| --- | --- | --- | --- | --- | --- | --- | --- |
| Agathidinae | *Alabagrus* sp1 | K | 8 | 7 | 1 | 2 | 18 |
| Agathidinae | *Alabagrus* sp2 | K | 0 | 2 | 0 | 0 | 2 |
| Agathidinae | *Bassus* sp1 | K | 7 | 2 | 1 | 0 | 10 |
| Agathidinae | *Bassus* sp2 | K | 4 | 0 | 0 | 0 | 4 |
| Agathidinae | *Bassus* sp3 | K | 10 | 13 | 2 | 0 | 25 |
| Agathidinae | *Bassus* sp4 | K | 3 | 0 | 1 | 0 | 4 |
| Agathidinae | *Pharpa* sp1 | K | 0 | 0 | 0 | 2 | 2 |
| Agathidinae | *Plesiocoelus* sp1 | K | 0 | 0 | 1 | 0 | 1 |
| Alysiinae | *Aphaereta* sp1 | K | 3 | 7 | 1 | 3 | 14 |
| Alysiinae | *Asobara* sp1 | K | 0 | 23 | 0 | 0 | 23 |
| Alysiinae | *Aspilota* sp1 | K | 0 | 0 | 1 | 2 | 3 |
| Alysiinae | *Aspilota* sp2 | K | 0 | 1 | 0 | 0 | 1 |
| Alysiinae | *Microcrasis* sp1 | K | 0 | 2 | 0 | 0 | 2 |
| Aphidiinae | *Aphidiinae* sp1 | K | 0 | 78 | 0 | 0 | 78 |
| Blacinae | *Blacus* sp1 | K | 0 | 0 | 0 | 5 | 5 |
| Blacinae | *Blacus* sp2 | K | 0 | 0 | 0 | 2 | 2 |
| Braconinae | *Bracon* sp1 | I | 25 | 5 | 1 | 0 | 31 |
| Braconinae | *Bracon* sp2 | I | 16 | 26 | 2 | 0 | 44 |
| Braconinae | *Bracon* sp3 | I | 1 | 2 | 1 | 0 | 4 |
| Braconinae | *Compsobraconoides* sp1 | I | 1 | 1 | 0 | 0 | 2 |
| Braconinae | *Habrobracon* sp1 | I | 0 | 3 | 1 | 0 | 4 |
| Braconinae | *Lapicida* sp1? | I | 0 | 2 | 0 | 3 | 5 |
| Braconinae | *Vipio* sp1 | I | 1 | 0 | 2 | 0 | 3 |
| Cardiochilinae | *Schoenlandella* sp1 | K | 0 | 17 | 0 | 0 | 17 |
| Cardiochilinae | sp1 | K | 0 | 1 | 1 | 1 | 3 |
| Cheloninae | *Chelonus* sp1 | K | 1 | 17 | 0 | 0 | 18 |
| Cheloninae | *Chelonus* sp2 | K | 0 | 10 | 4 | 3 | 17 |
| Cheloninae | *Chelonus* sp3 | K | 0 | 0 | 0 | 1 | 1 |
| Cheloninae | *Chelonus* sp4 | K | 2 | 9 | 0 | 0 | 11 |
| Cheloninae | *Chelonus* sp5 | K | 0 | 4 | 0 | 0 | 4 |
| Cheloninae | *Chelonus* sp6 | K | 0 | 4 | 0 | 0 | 4 |
| Cheloninae | *Chelonus* sp7 | K | 1 | 0 | 0 | 0 | 1 |
| Cheloninae | *Chelonus* sp8 | K | 3 | 0 | 0 | 0 | 3 |
| Cheloninae | *Chelonus* sp9 | K | 0 | 0 | 1 | 0 | 1 |
| Cheloninae | *Chelonus* sp10 | K | 0 | 0 | 1 | 0 | 1 |
| Cheloninae | *Chelonus* sp11 | K | 0 | 2 | 0 | 0 | 2 |
| Cheloninae | *Phanerotoma* sp1 | K | 3 | 12 | 0 | 0 | 15 |
| Cheloninae | *Phanerotoma* sp2 | K | 1 | 3 | 2 | 0 | 6 |
| Cheloninae | *Phanerotoma* sp3 | K | 0 | 0 | 0 | 1 | 1 |
| Cheloninae | *Phanerotoma* sp4 | K | 0 | 0 | 1 | 0 | 1 |
| Cheloninae | *Pseudophanerotoma* sp1 | K | 0 | 2 | 0 | 0 | 2 |
| Doryctinae | *Acrophasmus* sp1 | I | 7 | 1 | 1 | 0 | 9 |
| Doryctinae | *Curtisella* sp1 | I | 0 | 1 | 0 | 0 | 1 |
| Doryctinae | *Ecphylus* sp1 | I | 0 | 4 | 0 | 0 | 4 |
| Doryctinae | *Ecphylus* sp2 | I | 1 | 3 | 0 | 0 | 4 |
| Doryctinae | *Heterospilus* sp1 | I | 0 | 3 | 0 | 1 | 4 |
| Doryctinae | *Heterospilus* sp2 | I | 0 | 1 | 0 | 1 | 2 |
| Doryctinae | *Heterospilus* sp3 | I | 0 | 17 | 4 | 4 | 25 |
| Doryctinae | *Heterospilus* sp4 | I | 1 | 19 | 6 | 5 | 31 |
| Doryctinae | *Heterospilus* sp5 | I | 1 | 8 | 2 | 1 | 12 |
| Doryctinae | *Heterospilus* sp6 | I | 0 | 1 | 0 | 1 | 2 |
| Doryctinae | *Heterospilus* sp7 | I | 0 | 3 | 0 | 0 | 3 |
| Doryctinae | *Heterospilus* sp8 | I | 1 | 0 | 0 | 0 | 1 |
| Doryctinae | *Heterospilus* sp9 | I | 3 | 0 | 1 | 1 | 5 |
| Doryctinae | *Heterospilus* sp10 | I | 0 | 4 | 0 | 1 | 5 |
| Doryctinae | *Heterospilus* sp11 | I | 0 | 1 | 0 | 0 | 1 |
| Doryctinae | *Heterospilus* sp12 | I | 0 | 0 | 0 | 1 | 1 |
| Doryctinae | *Heterospilus* sp13 | I | 0 | 0 | 0 | 2 | 2 |
| Doryctinae | *Heterospilus* sp14 | I | 1 | 6 | 0 | 0 | 7 |
| Doryctinae | *Heterospilus* sp15 | I | 0 | 6 | 0 | 2 | 8 |
| Doryctinae | *Heterospilus* sp16 | I | 0 | 0 | 2 | 0 | 2 |
| Doryctinae | *Heterospilus* sp17 | I | 0 | 0 | 4 | 1 | 5 |
| Doryctinae | *Heterospilus* sp18 | I | 1 | 0 | 1 | 0 | 2 |
| Doryctinae | *Heterospilus* sp19 | I | 0 | 1 | 0 | 0 | 1 |
| Doryctinae | *Heterospilus* sp20 | I | 2 | 6 | 0 | 2 | 10 |
| Doryctinae | *Heterospilus* sp21 | I | 0 | 1 | 0 | 0 | 1 |
| Doryctinae | *Heterospilus* sp22 | I | 0 | 1 | 0 | 0 | 1 |
| Doryctinae | *Heterospilus* sp23 | I | 0 | 1 | 0 | 0 | 1 |
| Doryctinae | *Odontobracon* sp1 | I | 0 | 3 | 0 | 1 | 4 |
| Doryctinae | *Rhaconotus* sp1 | I | 1 | 2 | 4 | 0 | 7 |
| Doryctinae | *Rhaconotus* sp2 | I | 0 | 1 | 0 | 0 | 1 |
| Doryctinae | Sp1 | I | 0 | 1 | 0 | 0 | 1 |
| Euphorinae | *Aridelus* sp1 | K | 1 | 12 | 2 | 0 | 15 |
| Euphorinae | *Leiophron* sp1 | K | 0 | 10 | 0 | 0 | 10 |
| Euphorinae | *Leiophron* sp2 | K | 2 | 2 | 0 | 0 | 4 |
| Euphorinae | *Microctonus* sp1 | K | 0 | 0 | 0 | 2 | 2 |
| Euphorinae | *Microctonus* sp2 | K | 1 | 0 | 0 | 0 | 1 |
| Euphorinae | *Peristenus* sp1 | K | 0 | 1 | 0 | 0 | 1 |
| Homolobinae | *Exasticolus* sp1 | K | 0 | 1 | 1 | 0 | 2 |
| Homolobinae | *Homolobus* sp1 | K | 0 | 5 | 0 | 0 | 5 |
| Homolobinae | *Homolobus* sp2 | K | 0 | 0 | 1 | 0 | 1 |
| Hormiinae | *Cantharoctonus* sp1 | I | 0 | 1 | 0 | 0 | 1 |
| Hormiinae | *Hormius* sp1 | I | 1 | 1 | 1 | 0 | 3 |
| Hormiinae | *Hormius* sp2 | I | 5 | 23 | 0 | 0 | 28 |
| Hormiinae | *Hormius* sp3 | I | 0 | 1 | 0 | 1 | 2 |
| Hormiinae | *Hormius* sp4 | I | 3 | 33 | 0 | 0 | 36 |
| Hormiinae | *Oncophanes* sp1 | I | 0 | 1 | 0 | 0 | 1 |
| Hormiinae | *Oncophanes* sp2 | I | 2 | 2 | 0 | 0 | 4 |
| Hormiinae | *Oncophanes* sp3 | I | 1 | 5 | 0 | 1 | 7 |
| Hormiinae | *Oncophanes* sp4 | I | 0 | 4 | 0 | 2 | 6 |
| Hormiinae | *Pambolus* sp1 | I | 0 | 5 | 0 | 2 | 7 |
| Hormiinae | *Parahormius* sp1 | I | 1 | 1 | 0 | 0 | 2 |
| Hormiinae | *Rhysipolis* sp1 | I | 4 | 4 | 7 | 1 | 16 |
| Hormiinae | *Rhysipolis* sp2 | I | 0 | 1 | 0 | 1 | 2 |
| Hormiinae | *Rhysipolis* sp3 | I | 0 | 1 | 0 | 0 | 1 |
| Hormiinae | *Xenarcha* sp1 | I | 0 | 0 | 0 | 3 | 3 |
| Hormiinae | *Xenarcha* sp2 | I | 0 | 1 | 0 | 0 | 1 |
| Ichneutinae | *Oligoneurus* sp1 | K | 0 | 1 | 0 | 0 | 1 |
| Macrocentrinae | *Macrocentrus* sp1 | K | 1 | 0 | 0 | 0 | 1 |
| Mendesellinae | *Epsilogaster* sp1 | ? | 2 | 7 | 0 | 0 | 9 |
| Meteorinae | *Meteorus* sp1 | K | 1 | 0 | 1 | 0 | 2 |
| Microgastrinae | sp1 | K | 24 | 139 | 14 | 4 | 181 |
| Microgastrinae | sp2 | K | 5 | 0 | 4 | 0 | 9 |
| Microgastrinae | sp3 | K | 14 | 62 | 17 | 3 | 96 |
| Microgastrinae | sp4 | K | 1 | 0 | 2 | 0 | 3 |
| Microgastrinae | sp5 | K | 18 | 54 | 8 | 8 | 88 |
| Microgastrinae | sp6 | K | 26 | 47 | 8 | 2 | 83 |
| Microgastrinae | sp7 | K | 30 | 55 | 38 | 0 | 123 |
| Microgastrinae | sp8 | K | 11 | 55 | 7 | 1 | 74 |
| Microgastrinae | sp9 | K | 6 | 14 | 0 | 6 | 26 |
| Microgastrinae | sp10 | K | 65 | 37 | 2 | 0 | 104 |
| Microgastrinae | sp11 | K | 2 | 0 | 3 | 0 | 5 |
| Microgastrinae | sp12 | K | 3 | 6 | 3 | 0 | 12 |
| Microgastrinae | sp13 | K | 31 | 9 | 3 | 0 | 43 |
| Microgastrinae | sp14 | K | 1 | 6 | 2 | 0 | 9 |
| Microgastrinae | sp15 | K | 0 | 5 | 0 | 0 | 5 |
| Microgastrinae | sp16 | K | 6 | 43 | 5 | 12 | 66 |
| Microgastrinae | sp17 | K | 1 | 5 | 1 | 0 | 7 |
| Microgastrinae | sp18 | K | 0 | 2 | 1 | 2 | 5 |
| Microgastrinae | sp19 | K | 1 | 4 | 1 | 0 | 6 |
| Microgastrinae | sp20 | K | 2 | 12 | 1 | 0 | 15 |
| Microgastrinae | sp21 | K | 0 | 15 | 0 | 7 | 22 |
| Microgastrinae | sp22 | K | 0 | 0 | 0 | 1 | 1 |
| Microgastrinae | sp23 | K | 0 | 2 | 1 | 0 | 3 |
| Miracinae | *Mirax* sp1 | K | 4 | 17 | 0 | 6 | 27 |
| Opiinae | *Opius* sp1 | K | 30 | 81 | 2 | 27 | 140 |
| Opiinae | *Opius* sp2 | K | 0 | 3 | 0 | 0 | 3 |
| Opiinae | *Opius* sp3 | K | 0 | 2 | 0 | 0 | 2 |
| Opiinae | *Opius* sp4 | K | 4 | 4 | 0 | 0 | 8 |
| Opiinae | *Opius* sp5 | K | 1 | 10 | 1 | 8 | 20 |
| Opiinae | *Opius* sp6 | K | 11 | 37 | 2 | 3 | 53 |
| Opiinae | *Opius* sp7 | K | 1 | 6 | 1 | 1 | 9 |
| Orgilinae | *Orgilus* sp1 | K | 0 | 0 | 0 | 2 | 2 |
| Orgilinae | *Orgilus* sp2 | K | 4 | 0 | 0 | 0 | 4 |
| Orgilinae | *Stantonia* sp1 | K | 1 | 0 | 1 | 0 | 2 |
| Orgilinae | *Stantonia* sp2 | K | 0 | 4 | 3 | 2 | 9 |
| Rogadinae | *Aleiodes* sp1 | K | 1 | 0 | 0 | 1 | 2 |
| Rogadinae | *Aleiodes* sp2 | K | 0 | 4 | 0 | 0 | 4 |
| Rogadinae | *Stiropius* sp1 | K | 9 | 12 | 0 | 0 | 21 |
| Rogadinae | *Stiropius* sp2 | K | 0 | 7 | 0 | 0 | 7 |
|  |  |  | 447 | 1232 | 194 | 158 | 2031 |

